# Anisotropy of object nonrigidity: High-level perceptual consequences of cortical anisotropy

**DOI:** 10.1101/2024.09.10.612333

**Authors:** Akihito Maruya, Qasim Zaidi

**Affiliations:** Graduate Center for Vision Research, State University of New York, 33 West 42nd St, New York, NY 10036

**Keywords:** Object nonrigidity, cortical anisotropy, shape constancy, motion perception, cortical model, kinematic invariants

## Abstract

We demonstrate an unexpected anisotropy in perceived object non-rigidity, a higher-level perceptual phenomenon, and explain it by the population distribution of low-level neuronal properties in primary visual cortex. We measured the visual interpretation of two rigidly connected rotating circular rings. In videos where observers predominantly perceived rigidly-connected horizontally rotating rings, they predominantly perceived non-rigid independently wobbling rings if the video was rotated by 90°. Additionally, vertically rotating rings appeared narrower and longer than horizontally rotating counterparts. We decoded these perceived shape changes from V1 outputs incorporating documented cortical anisotropies in orientation selectivity: more cells and narrower tuning for the horizontal orientation than for vertical. Even when shapes were matched, the non-rigidity anisotropy persisted, suggesting uneven distributions of motion-direction mechanisms. When cortical anisotropies were incorporated into optic flow computations, the kinematic gradients (Divergence, Curl, Deformation) for vertical rotations aligned more with derived gradients for physical non-rigidity, while those for horizontal rotations aligned closer to rigidity. Our results reveal how high-level non-rigidity percepts can be shaped by hardwired cortical anisotropies. Cortical anisotropies are claimed to promote efficient encoding of statistical properties of natural images, but their surprising contribution to failures of shape constancy and object rigidity raise questions about their evolutionary function.

## Introduction

We present a remarkable instance of variations in a complex higher-level percept explained directly from the distribution of low-level neural properties in primary visual cortex by using a combination of mathematical derivations and computational simulations to reproduce the results of psychophysical experiments. The higher-level percept we demonstrate is an unexpected anisotropy in object nonrigidity, and the low-level properties are documented cortical anisotropies in numbers and tuning widths of orientation selective and direction selective neurons. The video in Figure 1A shows two connected Styrofoam rings on a rotating turntable. Most observers report that the two rings are rigidly joined and rotating together, although sometimes the top ring is reported as moving independently from the bottom ring and rolling or wobbling, seemingly defying physical plausibility. In the video in Figure 1B, the right pair of rings is almost always reported as moving independently, with some observers seeing more than just a slippage between the rings as a deformation in the rigid object consisting of the top ring wobbling relative to the bottom ring. The context reveals that one video is just a 90° rotation of the other (tilting the head by 90° also flips the percepts in each video), so clearly an explanation is needed for the increase in nonrigidity.

**Figure 1.**
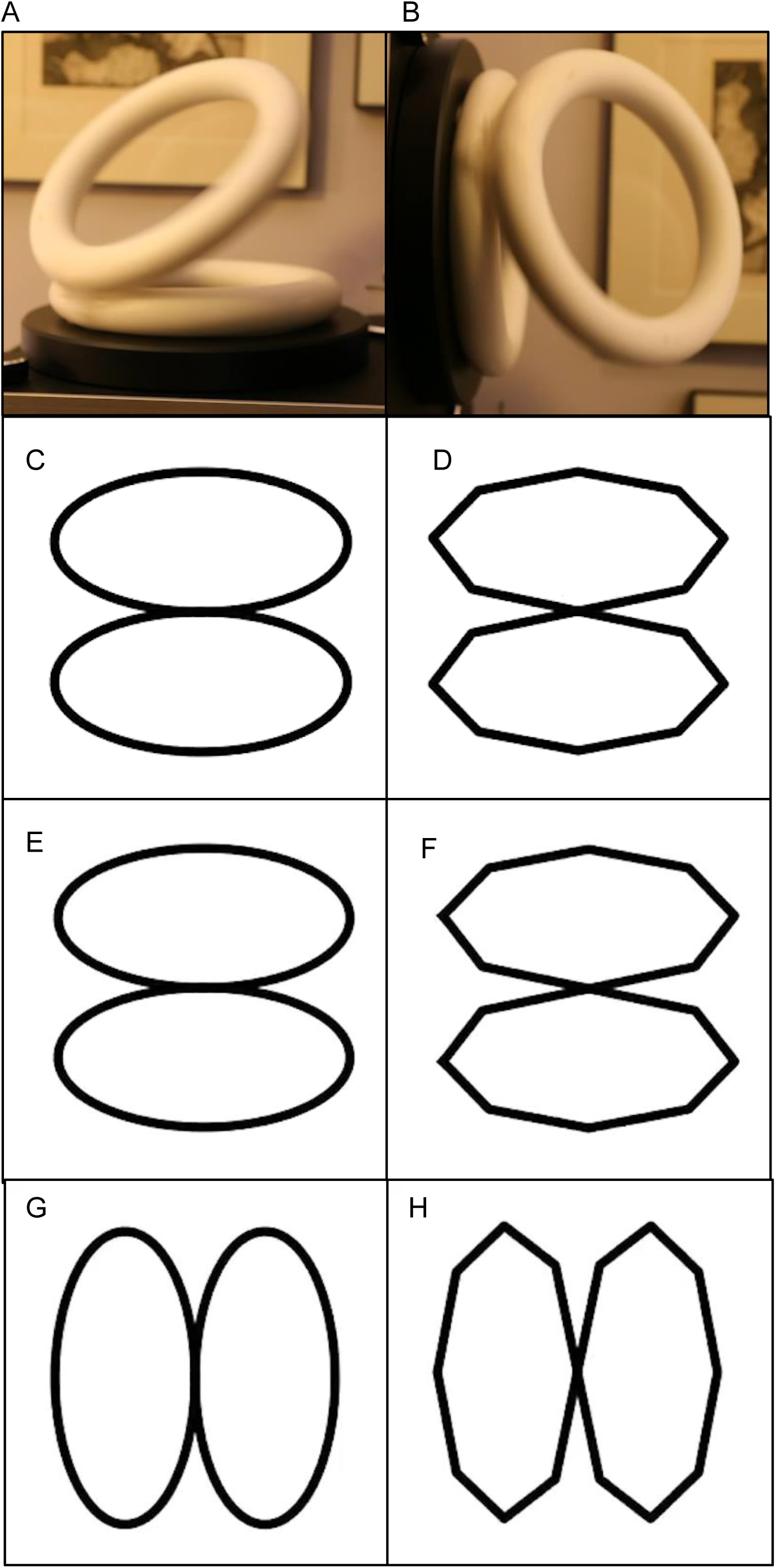
Anisotropy of a rotating ring illusion videos. A: Two Styrofoam rings glued together at an angle are seen to rotate together on the turntable. B: When rotated 90°, one ring appears to move independently and wobble against the ring on the turntable. C & D: Two circular (C) and octagonal (D) rings physically rotate together with a fixed connection at the junction. E & F: The two pairs of rings physically move independently and wobble against each other. For the circular rings, E is indistinguishable from C. However, with trackable vertices, F is discernable from D. G & H are the same as C & D except that they are rotated 90° and they both look nonrigidly connected.

3D-shape-from-motion models typically assume that objects are rigid, which simplifies the math^1^, but organisms generally change shape to move, eat, observe, mate, etc., so articulated and elastic motions are common in the real world^2^ and distinguish generically nonrigid moving organisms from generically rigid moving objects. The utility of this distinction is facilitated by human observers being as good at discerning 3D shapes of nonrigid objects from motion cues as they are for rigid objects^3^. Maruya and Zaidi (2024) showed that a rotating rigid object with two salient parts, such as the object in Fig 1, can appear rigid or nonrigid depending on speed of rotation and the presence of salient features^4^. At moderate to high speeds, the nonrigidity is predictable from the motion-energy flow which extracts motion vectors orthogonal to the contour, sometimes called the aperture problem^5^, whereas at slow speeds, feature tracking dominates^6,7^, if trackable features are present ^8,9,10,11,12,13^. The change in nonrigidity when the image is rotated by 90° could be due to anisotropy in shape or motion mechanisms. We investigate both as possible consequences of documented anisotropies in the population distribution of orientation and direction selective neurons.

To function successfully in the objective reality of the physical world, an organism’s perceptions must lead to accurate inferences of attributes such as shapes, sizes, locations, distances, motions, colors, textures, and kinds^14^. The earliest discovered cave drawings^15^ suggest that shape is the most important attribute for humans to identify objects and animals. In mathematics, the shape of a rigid object is a geometric attribute that remains unchanged across variations in location, rotation, and scale^16^, which corresponds to viewpoint invariance for a visual system. The shape of an object would not be a useful attribute if the mathematical definition was violated when an object was viewed from different angles. Unfortunately shape constancy is often violated. For example, horizontal parallelogram balconies on a tall building are seen as rectangles tilted up or down depending on viewpoint^17^. Such violations have been attributed to a tendency to assume right angles^17,18^, to try to maximize compactness or symmetry^19^, but the neural substrates of such processes are completely unknown. A more understandable kind of shape-constancy violation for 3D objects is caused by simply rotating the object in the frontal-plane. Cohen and Zaidi showed that the perceived depth of a wedge defined by shape-from-texture cues is a function of the orientation of the shape^20^. They linked this to a 2D anisotropy in the perception of an angle as a function of its orientation and showed that the 2D effect could be explained by a model based on the documented anisotropy in number and selectivity of orientation tuned cells in primary visual cortex of mammals which show the greatest concentration and narrower tuning for horizontal orientations, then vertical, and then oblique^21^ The anisotropy in primary visual cortex has been linked to encoding the statistical distribution of orientations in the world^22,23^, thus suggesting a common cause for the perceived effects of object orientation and the “oblique effect” where observers exhibit superior performance for stimuli presented along the cardinal (vertical and horizontal) axes as compared to oblique axes^24^. The cortical anisotropy has now been well documented for the ventral stream of primate brains^25^, where the anisotropy is a combination of radial and cardinal bias in V1 and mainly a cardinal bias in V4, raising the question of why evolution leads to brain properties that give more importance to the detection/discrimination of isolated orientations than to preserving perceived shapes across image rotations.

The neural bases of anisotropy of motion processing and perception is less well understood. Psychophysical anisotropies favoring cardinal directions over oblique directions have been shown for direction-discrimination but not for direction-detection of translating stimuli^26, 27, 28, 29, 30^. The anisotropies that have been documented in area MT^31, 32^ seem to not be systematic, although the number of cells measured is much less than in the V1 studies. There are some reports of direction selectivity for object motion in V4^33^, where there are documented anisotropies^25^, but the two issues have not been studied jointly. Given that most V1 simple cells receiving Magnocellular input from LGN are direction selective^34^, in our modeling, we assume that the direction preference and tuning of cells in mammalian striate cortex could be assumed to correspond to orientation preference along the axis orthogonal to the motion direction^21^. To our knowledge, anisotropies in motion based higher level perception, such as 3D-shape-from-motion or rigidity versus nonrigidity, have not been studied previously.

Shape and nonrigidity anisotropies are also easy to see when we switch from physical objects to computer graphics and orient the object to be symmetrical around the vertical or horizontal axis, such as the videos of the ring-pairs in Figures 1C – 1H. Figures 1C & 1D in the 2nd row are perspective projections of rigidly rotating configurations and Figures 1E & 1F in the 3rd row are perspective projections of rings that wobble and slide nonrigidly against each other. When attention is directed to the connecting joint, rigidity and nonrigidity are both easy to see in the octagonal rings because the vertices at the junction of the two rings either stay together or slide past each other. The circular rings, however, are without features, so rigidly rotating and wobbling rings generate identical retinal images and could be seen as rigidly rotating or nonrigidly wobbling, depending on how the images are processed by the visual system. If the image is rotated by 90 degrees, the nonrigidity becomes much more pronounced for both types of rings (Figures 1G & 1H in the 4^th^ row). Figures in the 2^nd^ and 4^th^ rows show a single frame of horizontally or vertically rotating rings, where the intersection of the two rings is in the center of the image. The vertically elongated rings appear noticeably narrower and longer than the horizontal ones. By turning the page 90 degrees, the reader can ascertain that this shape illusion is a function of object orientation.

Are the shape changes responsible for the increased nonrigidity, and is it due to a cortical anisotropy? We first show that these perceived shape changes can be decoded from V1 outputs, by considering anisotropies in the number and tuning widths of orientation-selective cells. Then we show empirically that when observers widened the vertically rotating ellipses or elongated the horizontally rotating ellipses so that the shapes matched, the perceived nonrigidity was still greater for the vertical rotation. We next incorporated cortical anisotropies into motion flow computations. The estimated motion fields were decomposed into velocity gradients of divergence, curl, and deformation. Rigidity of rotating 2D shapes can be tested by showing that only the curl component is non-zero^35^, but this is not true for the projections of the 3D rings, so we had to compare all three gradients generated by physical rotation versus physical wobbling. The gradients for vertical rotation of the rings more closely matched physical wobbling, while the gradients for horizontal rotation fell between physical wobbling and rotation, suggesting that hardwired cortical anisotropies can explain the increase in perceived nonrigidity from horizontal to vertical rotation.

## Results: Phenomenology, Modeling and Experiments

We will first examine the causes of the illusory static shape anisotropy and whether it can account for the dynamic nonrigidity anisotropy. If the shape illusion does not account for most of the motion illusion, we will then examine velocity anisotropy.

### Perceived Shape Anisotropy

Figures 2A & 2B present the two orientations of the rings in the phase that best demonstrates that the vertical rings look narrower and longer than the horizontal rings. This illusion is like anisotropy for an elongated diamond (Figure 2C & 2D). The diamonds have identical shape, but the vertical diamond looks narrower and longer. In Figure 2C & 2D, the angles *ϕ*_*h*_ and *ϕ*_*v*_ are physically equal but *ϕ*_*v*_ looks narrower (the subscript *h* indicates angles centered around the horizontal axis, while the subscript *v* indicates angles centered around the vertical axis). Similarly, *ψ*_*h*_ looks wider despite *ψ*_*h*_ and *ψ*_*v*_ being equal angles. As a result, the vertical diamond looks narrower. Cohen and Zaidi showed that flat 2D angles look broader when the central axis is obliquely oriented compared to vertical and modeled this as an effect of decoding the outputs of V1 cells, where those tuned to horizontal orientations were larger in number and narrower in tuning than those tuned to oblique orientations^20^. We now extend this to decoding vertical versus horizontally centered angles to explain the diamond illusion (Detailed explanation of computational procedures with equations in the Supplementary Section: Decoding Shape).

**Figure 2.**
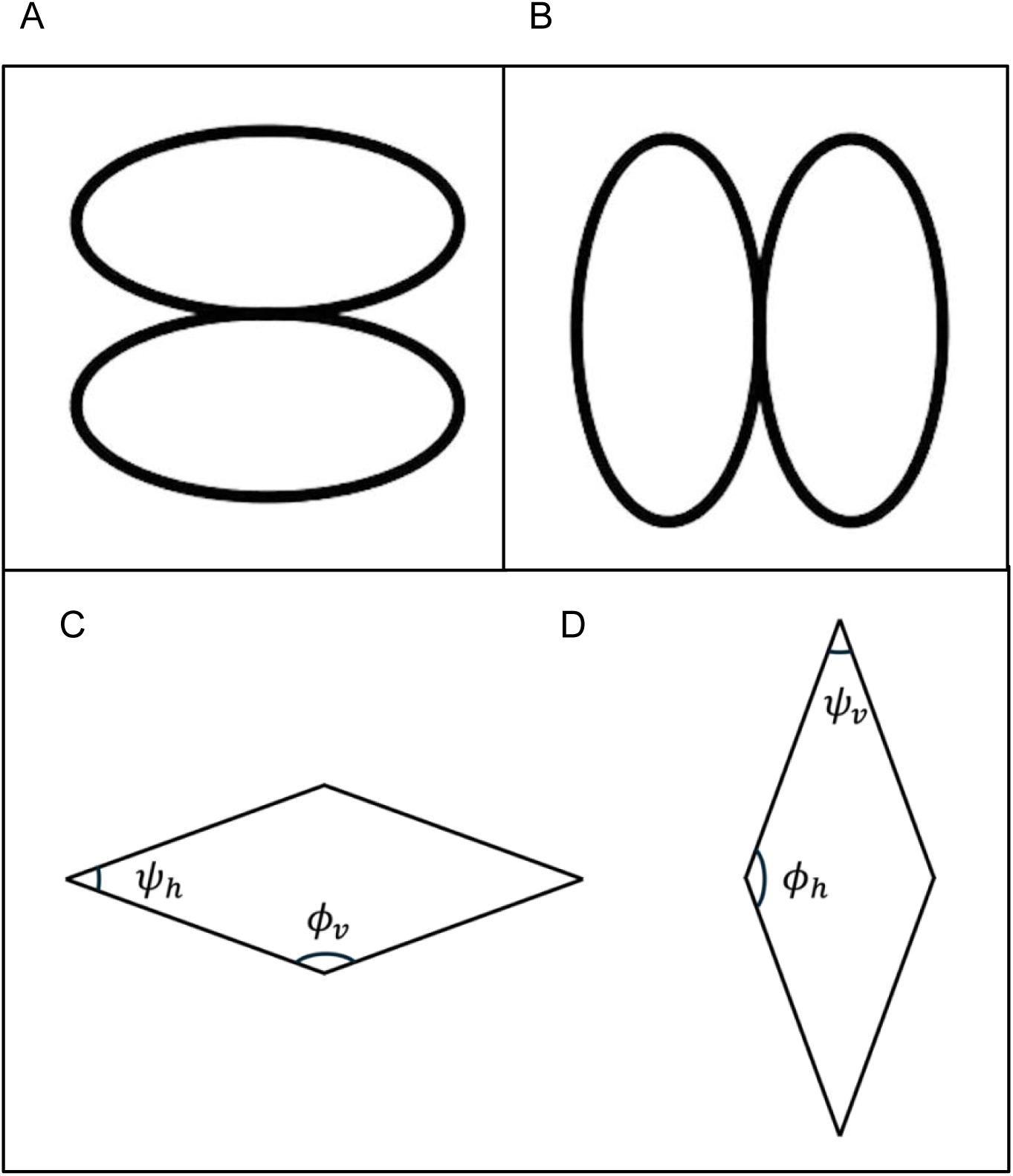
Anisotropy of the shape of the ring. A and B correspond to snapshots from Figure 1E and 1F respectively. Despite being physically identical shapes, B is perceived as vertically elongated and narrower in comparison to A. C and D: The shape illusion is similar for two elongated diamonds and can be explained by differences in perceived angles: *ϕh* is perceived to be wider than *ϕv* despite being physically equal, and *ψh* is perceived to be wider than *ψv*.

### Cortical Anisotropy of Orientation Selectivity

Figure 3A shows the orientation tuning widths of V1 simple cells we used in our model, centered at the preferred orientation, where 0° and ±180° represent horizontal preferred orientation, and ±90° represent vertical orientation, representing a summary of cat cortex measurements made with drifting gratings^21^. The heights of the curves reflect the relative number of cells tuned to each direction^21^. The left two panels in Figure 3B show the values of numbers and tuning widths plotted in terms of preferred orientation by averaging across opposite motion directions^21^. Cardinal orientations exhibit a higher number of cells and narrower tuning width. However, cells preferring vertical orientation are fewer and broader tuned than cells preferring horizontal orientation. There are more recent and more extensive data on primate V1 and V4^25^, but we did not use these because they provide only relative numbers, whereas Cohen & Zaidi^20^ found that tuning widths were more important than numbers, and our simulations reproduced that effect. The simulated orientation tuning curves use von Mises probability density functions to generate the tuning curve for all orientations over 360°, wider range than the 180°usual^36^ since we will be decoding angles where two orientations 180° apart are considered distinct. As a control, we also decoded angles from a cortex with isotropic distribution of orientation preference and tuning (Figure 3A).

**Figure 3.**
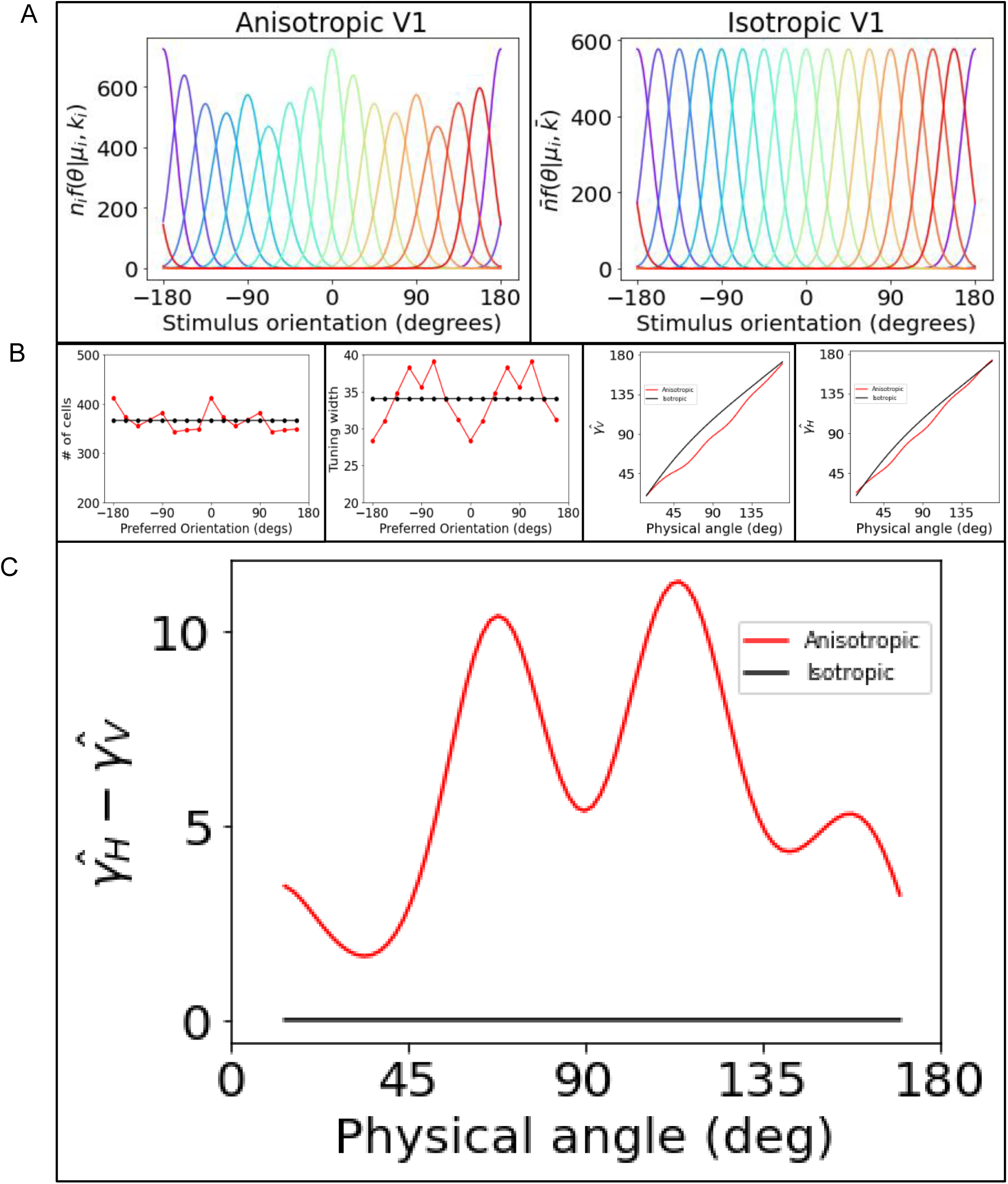
Cortical anisotropy. A: Orientation tuning curves of V1 simple cells were simulated by von Mises distributions to match the anisotropic and isotropic cortices and weighted by the number of cells (See text for references). B: Detailed parameters for the von Mises distributions are shown on the left two panels for the anisotropic (red) and isotropic (black) cortex. Decoded angles around the vertical and horizontal diverging axes respectively are shown on the right two panels. C: The decoded angle difference 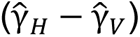 from the anisotropic cortex (red) and the isotropic cortex (dotted black). For the isotropic cortex, there is no difference in the decoded angles for γ_*v*_ and γ_*H*_. However, for the anisotropic cortex, γ_*H*_ is decoded to be broader than γ_*v*_.

### Perceived angle decoded from cortical responses

In Figure 2, angles around the horizontal axis (*γ_h_*) are perceived to be wider than physically equal angles around the vertical axis (*γ_v_*). We decoded the difference between them for angles ranging from 10-170° from an anisotropic cortex as well as an isotropic cortex. The left two panels in Figure 3B show the Li et al (2003) curves for simple cells plotted in terms of preferred orientation by averaging across opposite motion directions^21^. We assume that each angle is formed by two lines, to each of which we compute the responses of neurons of all preferred orientations, which are then normalized by a divisive gain control (see Supplementary Section: Decoding Shape). In primary visual cortex, neuronal responses are suppressed by the presence of surrounding stimuli. This suppression is orientation-specific, with the strongest suppression occurring when the stimuli in the receptive field and the surround share a similar orientation, and the weakest suppression occurring when they are orthogonal to each other^37,38,39^. The divisive gain control mechanism, involving self-normalization or dividing by the weighted sum of all other neurons (where the weight is highest for neurons with the same orientation preference), can achieve this neural interaction^40^, so the decoded orientation of a line also depends on the accompanying line’s orientation. The line orientations are decoded using vector sums of the normalized respones^41, 42^: In the left two panels of Figure 3B, the number of cells tuned to horizontal and vertical differ marginally, but the tuning width differences are clear with narrower horizontal tuning. The decoded orientations of the lines from the anisotropic cortex have a greater bias towards the vertical due to the broader orientation tuning of the vertically oriented RFs, being a more powerful factor than the larger number of cells tuned to the horizontal, but due to the divisive normalization creating a repulsion from the accompanying orientation, decoded orientations from both anisotropic and isotropic cortex will deviate from veridical.

Finally, the decoded angle is simply calculated as the difference between the two decoded line orientations. Right two panels in Figure 3B show decoded angles for the anisotropic cortex (red) and the isotropic cortex (black). Notice that the decoded angle for the isotropic cortex exhibits a slight overestimation due to divisive normalization, similar to the repulsion effect observed in the tilt illusion^42^. Figure 3C displays the decoded difference 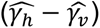 between an angle centered around the horizontal axis and a physically identical angle around the vertical axis, for the anisotropic cortex (red) and the isotropic cortex (black). Decoded responses from the isotropic cortex show no difference between the estimated horizontal and vertical angles. However, when decoded from the anisotropic cortex, angles around the horizontal axis γ_h_ are estimated to be wider than physically equal angles around the vertical axis γ_v_ which is a consequence of the vertical bias in decoded orientations of lines, and consistent with what is observed in Figure 2C & 2D. The difference varies with the magnitude of the angle but is greater than 5 degrees for all but very obtuse angles. We present this simulation to show that the illusory shape anisotropy in diamonds can be attributed to anisotropy in cortical populations. The estimated difference for a physical angle of 90° also explains why a square rotated 45° appears as a diamond which is elongated vertically^43^. Similar, but more complex decoding would yield a similar conclusion for the ellipses in Figure 2A & 2B. The qualitative inference is sufficient for our purposes, because the anisotropy assumptions from cat cortex are qualitatively similar but not quantitatively identical to measurements from primate cortex. Without the divisive normalization in Equation 3, the magnitude of the decoded difference is appreciably smaller. Cohen & Zaidi (2007) reproduced perceived differences in vertical versus oblique angles with maximum likelihood decoding by explicitly including cross-orientation suppression^20^.

It is worth comparing the diamond and ellipse anisotropy to horizontal and vertical differences that may relate to perspective. In possibly the most famous such illusion, Shepard’s table^44^, identical parallelograms appear very different in orthogonal orientations, when presented as parts of tables with legs in roughly perspective projection. The perspective context seems critical in this illusion, because the parallelograms compared without the context do not look as different across orientations, thus the illusion is unlikely to be governed by early cortical anisotropies. There is also a long history of failures to equate horizontal and vertical distances when perceiving three-dimensional scenes which have been explained without reference to neural mechanisms in terms of inappropriate application of depth scaling cues^45^. In addition, there are distortions in judgments of sizes of 3D objects as a function of their pose angle, especially the underestimation of sizes of objects pointing towards or away from the observer^46,47^. These perceptual distortions seem to be explainable in terms of back-transforms and misperceived ancillary cues, without needing cortical anisotropies.

### Effect of shape matching on perceived nonrigidity

Maruya & Zaidi (2024) showed that horizontally rotating elliptical rings with physical aspect ratios greater than 1.0 (narrower than circular rings) are perceived as more nonrigid than horizontally rotating elliptical rings with physical aspect ratios smaller than 1.0 (wider than circular), so the shape difference is a potential explanation for the nonrigidity anisotropy^4^. To test whether this explanation is sufficient, we had observers adjust shapes for vertical and horizontal rotations until they matched and then tested if matching projected shapes abolishes the nonrigidity anisotropy.

We presented pairs of vertically and horizontally rotating rings and asked 4 observers to match their shapes by stretching one of the pairs horizontally. Note that horizontal stretching makes the horizontally rotating rings narrower, and the vertically rotating rings wider, thus both counter the illusory percept of greater elongation for the vertically rotating rings. In Figure 4A, the 1st and 3rd rows show horizontal stretches of the 2D images ranging from 0% to 50%, while the 2nd and 4th rows illustrate 3D physical stretches of the rings, i.e. simulated stretches of the 3D rings observed as stretches of the projected 2D images of the rings. The differences between the image stretches and physical stretches become easier to see if enlarged to the size used in the experiment. Figures 4B and 4C present histograms of the horizontal stretch in the image domain (red) and the physical domain (blue) that match the shapes of horizontally rotating rings (left) and vertically rotating rings (right). To match shapes, observers stretched the horizontal rings and the vertical rings in the horizontal direction by approximately 20-30%. Individual results are shown in Supplementary Figure S1. Next, we presented selected pairs of rotating rings in random order and 4 observers reported which pair looked more independent and nonrigid by looking at the joint between each pair. We conducted four types of comparisons: Original, where two pairs of circular rings had identical dimensions; *H*_*i*_ & *H*_*p*_, where horizontally rotating rings were stretched to match the shape of vertically rotating rings, with the stretch applied either to the image (i) or physically before projection (p); *V*_*i*_ & *V*_*p*_, where vertically rotating rings were stretched to match the shape of horizontally rotating rings, with the stretch applied either to the image (i) or physically before projection (p); and Max, where both vertical and horizontal rings were maximally stretched in the image domain.

**Figure 4.**
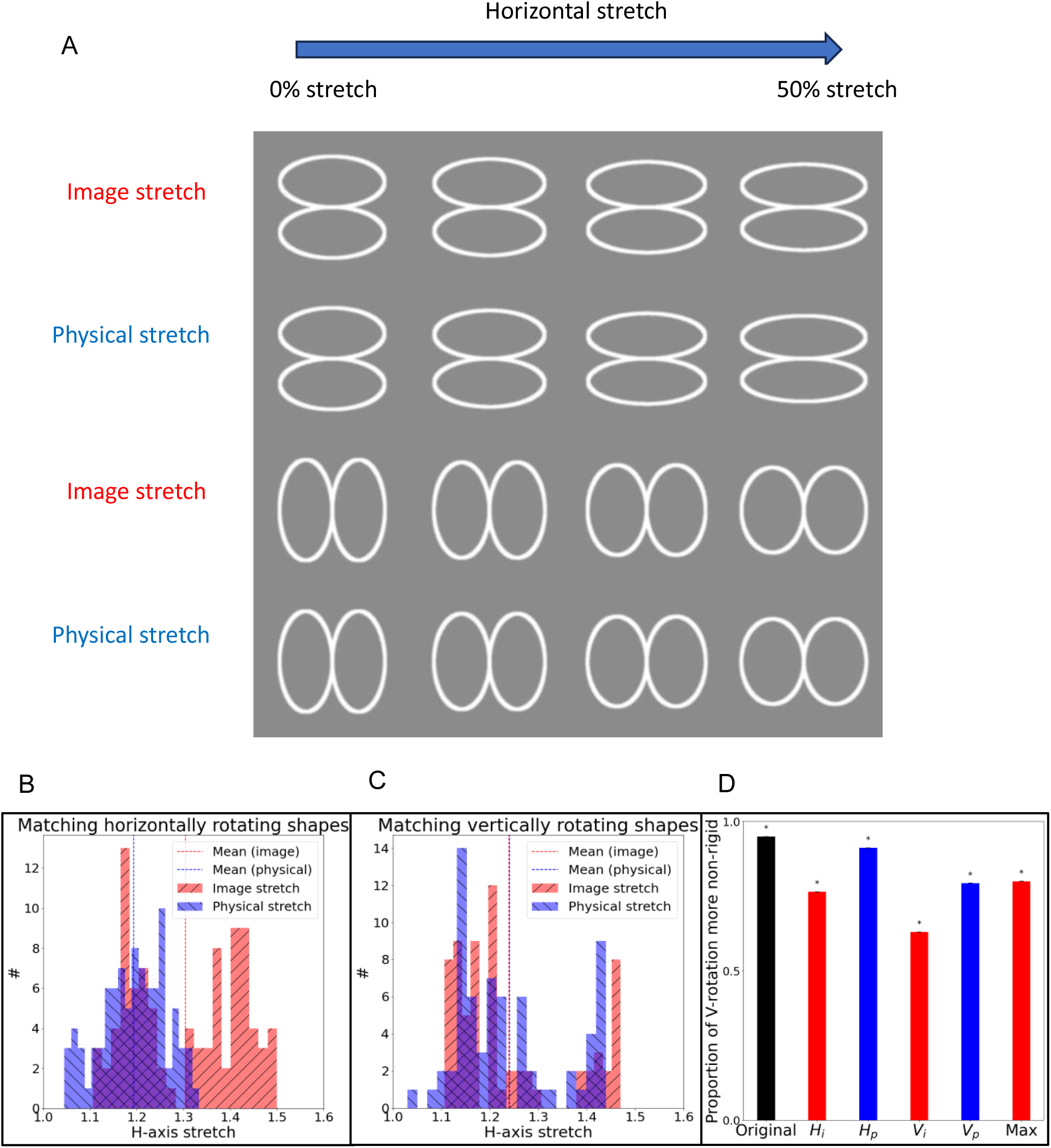
Effect of image and physical stretch on perceived nonrigidity. A: Examples of rings elongated to match the shape of orthogonally oriented circular rings. Subjects were allowed to stretch rings in image domain (first and third rows) or physically (second and fourth rows), ranging from 0% (left column) to 50% (right column). B: Histograms representing the extent of horizontal stretch in the image domain (red) and physical domain (blue) for vertically rotating rings adjusted to match the shape of physically circular horizontally rotating rings. C: Histograms representing the extent of horizontal stretch in the image domain (red) and physical domain (blue) for horizontally rotating rings adjusted to match the shape of physically circular vertically rotating rings. D: Probability of observers reporting vertically rotating rings as more nonrigid. Original: Two pairs of circular rings with identical dimensions. *H_i_* & *H_p_*: Horizontally rotating rings were stretched to match the shape of the vertically rotating rings, with the stretch applied either to the image (i) or physically before projection (p). *V_i_* & *V_p_*: Vertically rotating rings were stretched to match the shape of the horizontally rotating rings, with the stretch applied either to the image (i) or physically before projection (p). Max: Both vertical and horizontal rings were maximally stretched, as shown in the last column of Panel A, causing horizontally rotating rings to appear narrower and longer compared to vertically rotating rings.

Figure 4D shows the probability of observers reporting vertically rotating rings as more nonrigid. For the original pair, observers perceived vertically rotating rings as more nonrigid 95.0% of the time (black). Matching the shapes perceptually reduced the nonrigidity anisotropy by up to 40%, with a greater reduction observed in image-stretch (red bars) compared to physical stretch (blue bars). Individual results are shown in Supplementary Figure S2. In an ancillary forced choice experiment, observers picked the image stretch as a better shape match than the physical stretch (61%), possibly because the shape anisotropy is an illusory stretch of the image. However, the nonrigidity anisotropy remains significantly above chance levels despite the shape matches so the perceived shape does not account for almost 60% of the effect. As a critical test, we took the most stretched horizontal and vertical rings (Figure 4A Rightmost rings in 2^nd^ and 4^th^ row) and asked 4 observers to choose which pair was more nonrigid when rotated. Despite the horizontal rings being much narrower than vertical rings, the opposite of the original condition, observers picked the vertical rings as more nonrigid on 80% of the comparisons. Therefore, the difference in percepts of rigidity between horizontally and vertically oriented rotating pairs of rings cannot be accounted for solely by the effects of cortical anisotropy on perception of angles, and hence shapes, in vertically- and horizontally-oriented stimuli. opening the possibility that anisotropy of low-level motion detectors also plays a role.

### Nonrigidity anisotropy from cortical anisotropy in direction selectivity

The cat cortical anisotropy was measured with drifting gratings^21^, so we make the reasonable assumption that the anisotropic distribution of numbers and tuning widths also represents direction selective cells. To simulate the effect of spatial anisotropy on optic flow, we computed local motion energy for each pixel of the videos (Detailed explanation of computational procedures with equations in the Supplementary Section: DECODING NON-RIGIDITY VERSUS RIGIDITY). Each motion-energy unit consisted of a quadrature pair of 3D Gabor filters in the standard form ^48–55^. We simulated 8 spatial orientations from 0 to 7/8π, 5 temporal orientations from −π/4 to π/4) and spatial frequencies from 0.28 and 0.14 cycles/pixel. For the anisotropic cortex, the half-height orientation bandwidths were matched to cat cortex estimates^21^ using a standard equation^56^.

Each filter was convolved with the video. The responses of each quadrature pair were squared and summed to produce a phase-independent response, a well-known method^48–55^. Figure 5 A and B illustrate the direction tuning curves for isotropic and anisotropic cortices, weighted by the number of cells. In the anisotropic cortex, cells that prefer a vertically moving bar (either upwards or downwards) exhibit narrower tuning and are more numerous compared to cells that prefer oblique or horizontally moving bars.

**Figure 5.**
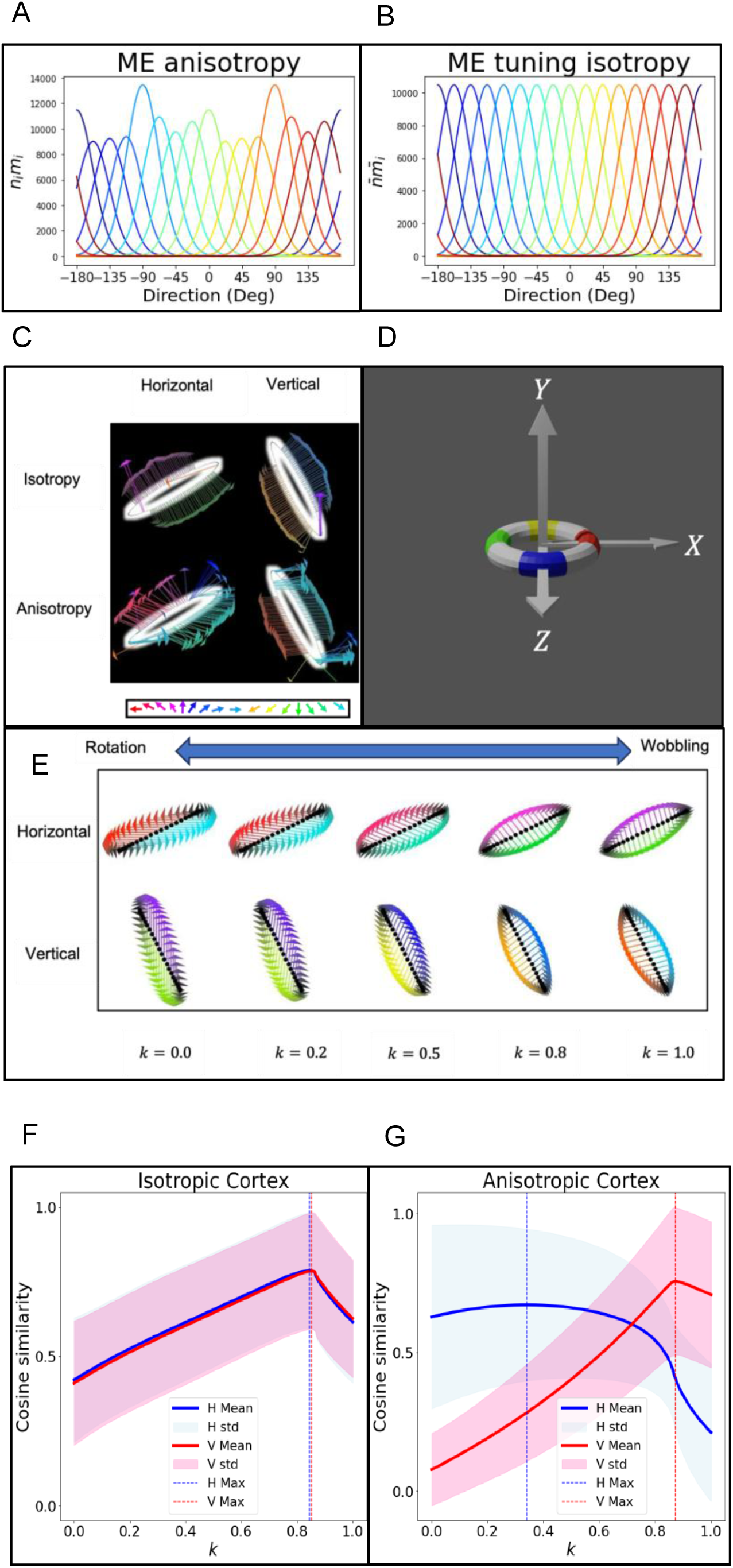
Comparing optic flows from isotropic cortex and anisotropic cortex to templates for rotation and wobbling. A: Tuning curves for direction selective cells reflecting documented anisotropies in width and number of cells. B: Tuning curves for direction selective cells for an isotropic corticex. C: Optic flow fields generated by isotropic cortex (top row) and anisotropic cortex (bottom row) for physical horizontal rotation (left column) and vertical rotation (right column). Anisotropic cortex generates vectors pointing horizontally. D: Depiction of rotation and wobbling axes. E: Physical velocity field for horizontal (top) and vertical (bottom) rotations mixed with physical wobbling (Weights k=0-1). F & G Cosine similarity of ME optic flows from isotropic (F) and anisotropic (G) cortex to different templates (*k*=0: physical rotation, *k*=1: physical wobbling, shown in Fig. S6 (**a**)). Isotropic cortex leads to no difference between best fitting *k* for horizontal (blue) and vertical (red) rotation. However, cortical anisotropy results in different best fitting *k*: more wobbling for vertical rotation.

To obtain the best estimate of velocity per point using the whole family of filters, we computed optic flow by estimating the motion vector that minimized the difference between the predicted and measured motion energies^57^. Figure 5C shows optic flow fields for different conditions of rotation of a single ring, horizontal rotation ℝ_*y*_ around the y-axis (left column), and vertical rotation ℝ_*x*_ around the x-axis (right column), for the isotropic cortex (top row), and the anisotropic cortex (bottom row). The color of the vector corresponds to the direction and the length represents the magnitude of the estimated velocity. Most of the optic flow vectors from the isotropic cortex are orthogonal to the contour. In contrast, for the anisotropic cortex, the optic flow vectors shift towards the horizontal direction, as indicated by the red and turquoise colors.

To understand what this horizontal shift in the optic flow implies for perceived rotation versus wobble (which adds rotation around the z-axis), we derived the velocity field for different combinations of physical rotation and wobbling of a single ring, with the wobbling weight *k* ranging from *k* = 0∼*rotation* to *k* = 1∼*wobbling*. We began with the vector which defines the position of every point on a ring lying on the *X* − *Z* plane for each angular coordinate of the circular ring (Figure 5D), and calculated the projection for every combination of rotation and wobble for a given angular velocity and tilt of the ring. Then, the velocity field was calculated by the partial derivative with respect to time.

Figure 5E illustrates the velocity fields for horizontal rotation (top) and vertical (bottom), obtained with wobbling weights (*k*) increasing from left to right, from pure rotation to pure wobbling. For horizontal rotation (top row), horizontal vectors are predominant when *k* = 0 and vertical vectors at *k* = 1, with the opposite for the vertical rotation (bottom row). These velocity fields serve as templates, and the cosine similarity between these derived templates and the estimated optic flows in Figure 5C is calculated at each point and time for both the isotropic and anisotropic cortex (Figures 5F & 5G) and then averaged over all points for a complete cycle. The blue trace represents mean cosine similarity for horizontal rotation, and the red trace for vertical rotation. As expected, for flows calculated from an isotropic cortex, cosine similarities are the same for horizontal and vertical rotations with the best fit at *k* = 0.84. For flows from the anisotropic cortex, horizontal and vertical rotations result in different cosine similarities. The horizontal rotation cosine similarity curve is relatively flat from *k* values 0.0 to 0.7 with a peak around 0.4, while the vertical rotation curve is steep and peaks at a higher *k*, close to 0.9. Thus, if generated velocity fields were compared to velocity field templates for physical rotation and wobbling, it would, explain why observers perceive more nonrigidity when the two rings rotate vertically. However, having all possible templates is biologically implausible, so next we explore how a more biologically plausible factoring of the velocity fields can lead to similar comparisons that predict the nonrigidity anisotropy.

### Nonrigidity anisotropy from biologically plausible filters for differential invariants

In kinematics, which describes the motion of objects without considering the forces that cause them to move, motion fields are analyzed in terms of the local differential invariants, divergence, curl, and deformation after excluding translation parallel to the image plane, and these analyses have been useful in understanding object motion from optic flow^35,58,59,60^. For the single ring with some combination of rotation and wobble, the divergence field is perpendicular to the contour and captures expansion/contraction of the projection (Figure 6A); the curl field is tangential to the contour and captures rotation; the deformation fields captures shear caused by the difference between orthogonal axes in contraction/expansion or shear and can be combined into one measure giving direction and magnitude of deformation^60^. Besides providing understandable summary measures, these operators also have biological validity. Zhang et al. (1993) demonstrated that position-independent detectors of curl and divergence can emerge through simple Hebbian learning, from MT-like receptors to MST-like receptors^61^. Physiological studies on nonhuman primates and imaging studies on humans have identified neurons in areas MST, STPa, and STS that preferentially respond to the curl and div components of optic flow^62, 63, 64, 65^. In addition, neural activity sensitive to def has been found in dorsal MST^66^, and human psychophysical experiments have demonstrated that perceived slant is proportional to def ^58, 67, 68^.

**Figure 6.**
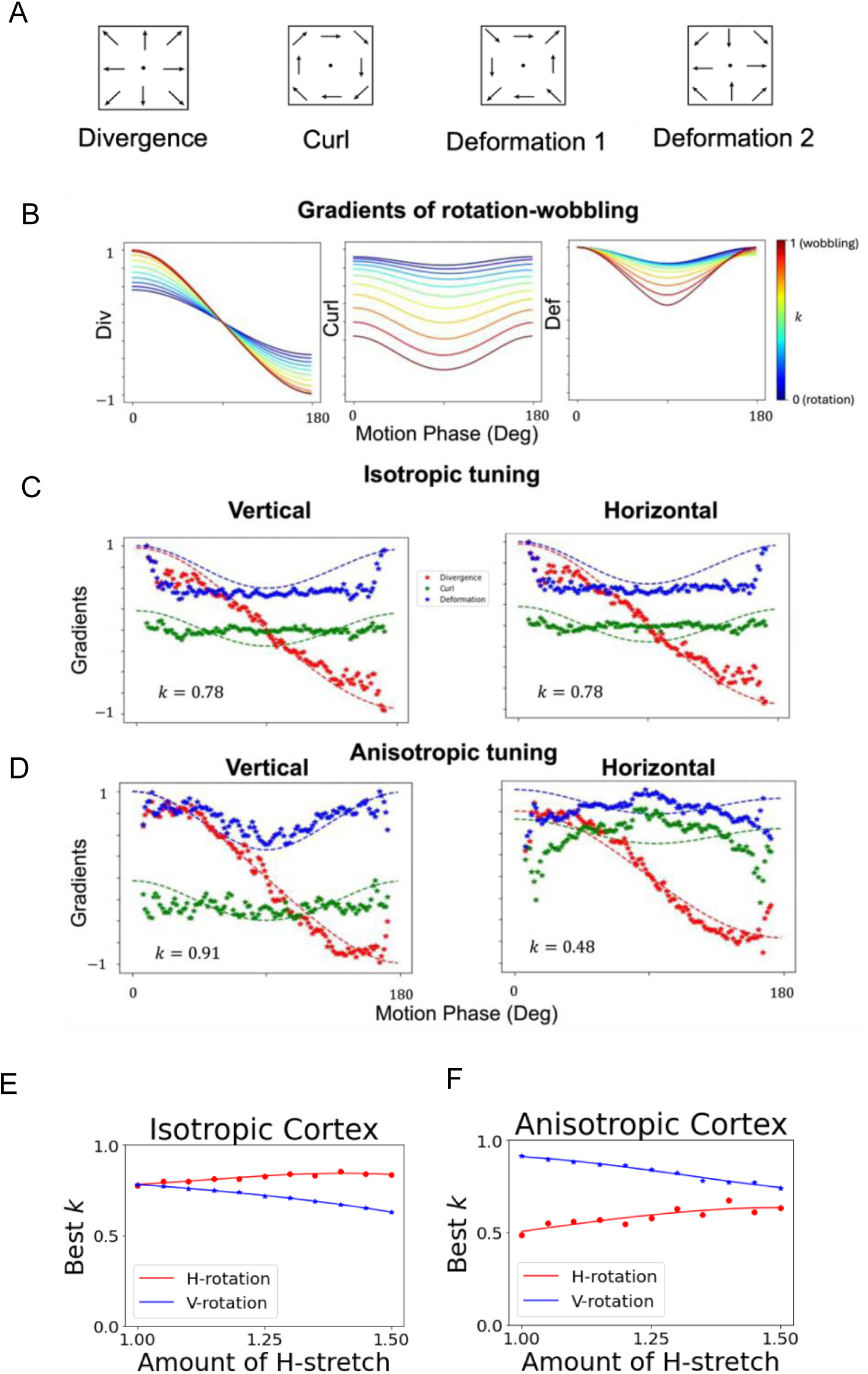
Gradients of ME flow from the rotating ring align with percepts. A: Examples of differential invariants. B: Gradients of rotation to wobbling, divergence (left), curl (middle), and deformatiosn1 (right) as a function of rotational phase (0°-180°: half a cycle). These gradients illustrate that with increasing emphasis on wobbling from k=0 (blue curves) to k=1 (red curves), variability increases across all three gradients, while rotational fields consistently exhibit higher values of curl and deformation. C: Isotropic Tuning Gradients for Vertical (left) and Horizontal (right) Rotation: Similar gradients are observed for both orientations, with optimal fitting by curves at a wobbling weight of k=0.78. D: Upon adjusting filter numbers and tuning widths to match V1 anisotropies before velocity field calculations, gradients for vertical rotation closely correspond to physical wobbling (k=0.91), while those for horizontal rotation indicate lower levels of wobbling (k=0.48). E & F : Best *k* as a function of image stretch. E for isotropic cortex and F for anisotropic cortex. A greater amount of stretch leads to increased wobbling during horizontal rotation (red) and reduced wobbling during vertical rotation (blue). For an anisotropic cortex, the higher amount of wobbling during vertical rotation is maintained, consistent with experimental results. Furthermore, at maximum stretch, the isotropic and anisotropic cortices predict different outcomes: the isotropic cortex suggests that horizontal rotation should be perceived as more nonrigid, while the anisotropic cortex suggests that vertical rotation should appear more nonrigid.

To derive divergence, curl, and deformation fields for physical rotation and physical wobbling, the dot product of the velocity fields derived in Equation S17 as a function of *k* (illustrated in Figure 5E) was taken with each operator corresponding to those illustrated in Figure 6A, and then integrated across space to obtain one number for each invariant per phase of motion (Detailed explanation of computational procedures with equations in the Supplementary Section: Nonrigidity anisotropy from biologically plausible filters for differential invariants)

Figure 6B displays the values of derived integrated gradients for divergence, curl, and deformation from left to right as a function of the motion phase. The colors represent different wobbling weights (*k*), with warmer colors indicating more wobbling and cooler colors representing more rotation. For all three gradients, the velocity field for rotation exhibits lower variance across different motion phases because the local velocity vectors consistently point in horizontal directions, so changes in the value result solely from alterations in shape due to projection. The wobbling field exhibits higher variance because the projected shapes and thus the local velocities change more over time. The value of Divergence varies from positive to negative representing divergence versus convergence, and the amplitude is greater for wobble because of the additional axis of rotation. Curl for the rotational velocity field is high, but almost constant, reflecting the minimal change in projected shape, whereas the greater projected shape changes for wobbling lead to smaller values of curl but a larger amplitude of variation. The derived deformation estimates have a similar variation. Taken together, the three derived gradients for the physical motions provide templates for distinguishing between relative weights of rotation and wobble, so next we estimated the values of the gradients for the optic flow fields for the single rotating ring computed from the isotropic and anisotropic cortex (Figure 5C). For cortical isotropy (Figure 6C), the divergence, curl, and deformation are identical for vertical and horizontal orientation (red symbols represent divergence, green represent curl, blue represent deformation). The dashed smooth curve in the corresponding color indicates the best fit of the derived physical motion. For both orientations, the best fits occur at a wobbling weight of *k* = 0.78, consistent with the computational results in Maruya and Zaidi ^4^ using CNNs that showed that motion-energy flows support percepts of wobble, because they predominantly signal motion orthogonal to contours. The cortical anisotropy distorts the optic flow so that the optimal fit for vertical rotation (*k* = 0.91) corresponds closer to physical wobbling than the fit (*k* = 0.48) for horizontal rotation. The analysis in terms of differential invariants thus provides an explanation for the anisotropy of nonrigidity that matches the one provided by template matching.

To further test the efficacy of this model, we tested if it could predict reduced anisotropy due to horizontal shape stretching in the shape matching experiment. We took the videos of the rotating single ring and applied horizontal image stretches ranging from 0% to 50%, and calculated the optic flows generated by the isotropic and anisotropic cortex similar to that shown in Figure 5C. Then we calculated the three differential invariants at each image stretch and estimated the optimal *k* as in Figures 6C and 6D for horizontal rotation and vertical rotation. For both isotropic (Figure 6E) and anisotropic (Figure 6F) cortices, increased stretching during horizontal rotation (red) results in a higher amount of wobbling, while increased stretching during vertical rotation (blue) leads to a reduced amount of wobbling, consistent with the experimental result that reduced anisotropy is observed with a greater amount of horizontal stretch.

However, for the anisotropic cortex vertical rotation maintains higher nonrigidity across different amounts of stretch, which aligns with the results in Figure 4D, while the isotropic cortex predicts the opposite. Moreover, the isotropic cortex simulations predict least anisotropy at 0% stretch and most at 50%, which is the opposite of the experimental results, while the anisotropic predicts that the anisotropy reduces with stretch but does not disappear, consistent with the experimental results. Taken together, the simulations from the anisotropic cortex explain the experimental measurements of non-rigidity.

## Discussion

Despite the abundance of nonrigid 3D objects and organisms in the world, and despite the evolutionary advantage of being able to judge their actions from shape deformations, nonrigid motion perception is an understudied phenomenon^3, 69, 70–78^. Maruya and Zaidi^4^ showed that the horizontal rotation of two rigidly connected circular 3D rings can appear either rigid or nonrigid (rings wobbling and rolling independently) if the rotational speed is sufficient to activate motion-energy cells. They found that the nonrigid perception arises from populations of motion energy cells signaling motion orthogonal to the contour. The nonrigidity can be decreased and even abolished if the rings contain salient features that can be used for feature-tracking and if the shape does not support a prior for spinning motion^4^. We strengthen this link by showing that the perceived nonrigidity is strongly affected by the cortical anisotropy for motion-direction selective neurons.

The nonrigidity anisotropy we demonstrate is as interesting as it is unexpected. We show that the hard-wired cortical anisotropy is powerful enough in modifying the optic flow from the vertically rotating rings towards the horizontal directions to counter the effects of salient features that evoke feature-tracking and overcome the effects of shape-based priors in the horizontal rotation (Video in Figure 1H).

A model that explains the complete percept of the 3D rings is beyond the scope of this paper, so we concentrate on the perception of nonrigidity versus rigidity and find surprisingly that variations in this high-level percept can be explained by anisotropies in the neural population, possibly even at the primary visual cortex. That certain variations in a high-level percept such as rigidity are a function of hardwired low-level neural properties is remarkable. Other than Maxwell’s success in identifying cone photoreceptors from color matches, there are almost always too many degrees of freedom to model anything definitive about neural mechanisms from psychophysical results. In addition, it seems that there is little information about higher-level neural processes in lower areas^79^. We may have succeeded because we picked a single higher-level aspect of a percept and a well-documented property of visual cortex, so our strategy may be worth repeating.

To link motion-energy to anisotropy we found a combination of computer simulations and mathematical derivations to be extremely useful. For calculating the optic flow for rings rotating horizontally and vertically we used banks of motion-energy filters, and adjusted their spatial envelopes to compare the effects of V1 cortical anisotropy to isotropy. To understand the physical motions that best describe the optic flows, we mathematically derived the velocity fields of rings undergoing different weighted combinations of rotation and wobbling motion (given by the parameter *k*), and correlated them with the optic flows. For optic flows generated from an isotropic cortex, the best fitted weights were the same for horizontal and vertical rotation. However, the change in the optic flow caused by an anisotropic cortex predicted different physical combinations for horizontal and vertical rotations. Vertical rotation aligned closely with physical wobbling, while horizontal rotation aligned closer to physical rotation, matching the perceptions reported by our observers. We realized that correlating different templates for different motions to the optic flow from an object is computationally expensive and biologically implausible, not to mention that the templates would not generalize across object shapes. Therefore, we explored a more efficient and plausible model using differential invariants.

Differential invariants from kinematics have the advantage of not being restricted to precise shapes, similar to direction-selective filters being more general than matching specific features across frames, and they can be calculated with combinations of biologically plausible filters. Consequently, we derived the differential invariants mathematically for the physical combinations of rotation and wobbling and plotted their values as a function of motion phase. Then we computed the values of the differential invariants given by the optic flows obtained from isotropic and anisotropic cortex and compared them to the values for different combinations of rotation and wobbling. The results strengthen our contention that the anisotropy in nonrigidity arises from the hardwired cortical anisotropy in numbers and tuning widths of direction selective neurons. From our previous work, we learned that the optic flows from motion-energy units can lead to seeing rigid objects as nonrigid, while the presence of trackable features increases the probability of rigid percepts^4^. Figure 1H demonstrates that the cortical anisotropy biases the optic flow enough to overcome the rigidity signals provided by trackable features.

An interesting corroboration of our model is provided by viewing the videos in Fig 1 in the peripheral visual field. The well-established centrifugal bias for motion in the periphery^80^ implies that in the visual periphery below a central fixation point, there will be a bias towards vertical motion, and in the visual periphery left of a central fixation point, there will be a bias towards horizontal motion. Based on these biases, our model predicts that the horizontal rotation should appear more rigid in the left visual field than on the lower, and the vertical rotation should appear more rigid in the lower visual field than on the left. For the two authors, the prediction is qualitatively correct, as the horizontal rotation shifts from rigid to non-rigid from left to lower. The vertical rotation appears more rigid in the lower visual field than on the left as predicted, but both still look non-rigid, indicating that the centrifugal bias cannot overcome the anisotropy in the distribution of orientation preferences and tuning-width that we use in the modeling.

To summarize the main points, our study demonstrates that vertically rotating rings appear more nonrigid than horizontally rotating rings. The shape of the vertically rotating rings seems narrower and longer compared to the horizontally rotating rings. By matching the shapes, observers were able to reduce perceived anisotropy by up to 30%, yet vertically rotating rings were still perceived as more nonrigid after shapes were matched. We incorporated anisotropy into our motion-energy computations and found that the extracted optic flow patterns from anisotropic cortex align with templates for physical wobbling for the vertically rotated rings but not for the horizontally rotated rings. Matching to derived differential invariants as a biologically plausible model, strengthened our conclusions.

In this study, an unexpected high-level perceptual phenomenon is explained purely by consideration of the numbers and qualities of response properties of cells in the mammalian visual cortex. This goes far beyond simply showing that the responses of particular cells correlate with particular aspects of a percept or that stimulating particular cells can change a percept in a particular fashion. Here, known non-uniformities in the numbers of cells sensitive to different stimuli and the qualities of those sensitivities affect estimates of the perceptual invariants from which we judge the three-dimensional motion of objects in the world. Understanding how such anisotropies in cellular responses, when those responses are combined over populations of cells, give rise to distinct percepts goes well beyond previous linkages between cellular and mental events. We hope that our experiments and models encourage more psychophysical and neurophysiological investigation of links between low-level hard-wired cortical properties and high-level perceptual inferences.

## Experimental methods

### Stimulus generation

Using Python, we created two circular rings connected rigidly at a 60 degree angle, rotating around either a vertical axis or a horizontal axis oblique to both rings. The joint’s speed when passing fronto-parallel was 6°/s, with a ring diameter of 3 degrees of visual angle (dva), resulting in an angular speed of 0.3 cycles per second (cps). We presented pairs of vertically and horizontally rotating rings and asked observers to match their shapes. The rings could be stretched horizontally from 0% to 50%, either in the image domain or the physical domain. Horizontal stretches in the image domain transformed the projected ellipses for the horizontal rotation into narrower and longer ellipses, and into wider and shorter ellipses for the vertically rotating axes. In the physical domain, stretches transformed the circular rings into elongated ellipses, and transformations in the projected images were similar to those for the image stretches. Figure 4A shows examples of elongated rings as a function of horizontal stretch in the image domain (first and third rows) and the physical domain (second and fourth rows). The matching was done while the rings were rotating in the same phase.

### Stimulus display

The videos were displayed on a VIEWPixx/3D (VPixx Technologies, Saint-Bruno-de-Montarville, QC, Canada) at 120 Hz. PsychoPy was used to display the stimulus and run the experiment. The data were analyzed using Python. An observer’s viewing position was fixed with a chinrest to ensure the video was viewed at the same elevation as the camera position. The chinrest was 1.0 meters from the screen which subtended 30 by 17 dva. The centers of the rings were 1.6 dva to the right and left of the center of the screen.

### Psychophysical procedures

Observers were asked to use the mouse to stretch the target shape to match the test shape. After three settings, the median stretch was taken, and each matched shape in the image and physical domains was presented side by side for the observer to select which of the two methods generated the shape most like the orthogonally oriented original. During the shape matches, observers were allowed to move their eyes freely across the two shapes. After the best shape matches were determined, we presented a vertically rotating ring pair and a horizontally rotating ring pair side by side, and observers were asked to look at the joint between the rings in each pair and select the more nonrigidly moving pair. For judging nonrigidity, observers were instructed to attend to the joint between the two rings in each pair. There were 5 comparison conditions (Figure 4D): Original: Two pairs of circular rings of identical dimensions; H_i_ & H_p_ indicate that the horizontally rotating rings were stretched to match the vertically rotating shape and V_i_ & V_p_ indicate that vertically rotating rings were stretched to match the horizontally rotating shape, with the stretch applied either to the image (i) or physically before projection (p). Max: Both vertical and horizontal rings were stretched maximally like in the last column of Panel A, so horizontally rotating rings were perceived as narrower and longer than vertically rotating rings. The set of conditions was repeated 40 times. Measurements were made by four observers with normal or corrected-to-normal vision. Observers gave written informed consent. All experiments were conducted in compliance with a protocol approved by the institutional review board at SUNY College of Optometry, adhering to the tenets of the Declaration of Helsinki.

## Author contributions

AM and QZ designed the study. AM programmed and ran the experiments. AM & QZ analyzed and modeled the results. AM & QZ wrote the paper.

## Acknowledgements

NEI grants EY035085 & EY035838

## Resource availability

All data and software will be publicly available on digital depositories.

## Data availability

All data and code are available in the following GitHub repository: https://github.com/amaruya/Anisotropy-of-object-nonrigidity-High-level-perceptual-consequences-of-cortical-anisotropy/tree/main

## information

We present the formal version of the computational and analytic derivations described in the text (including equations), followed by a table describing all the symbols. This section also includes figures for individual observers.

### STIMULUS GENERATION & PROJECTION

We created 3D circular, rotating/wobbling rings by applying the equations for rotation along the *X*, *Y*, and *Z* axes in a 3D space to all points on the rendered object. The rotational matrix around each axis is given by:

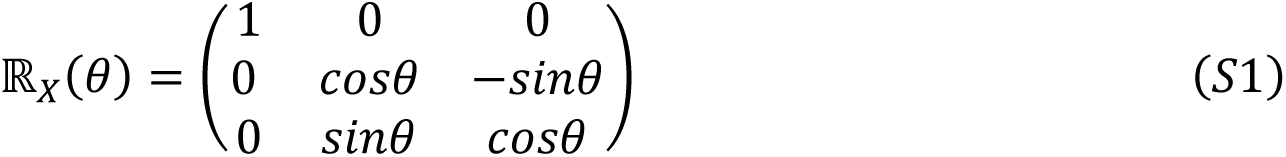

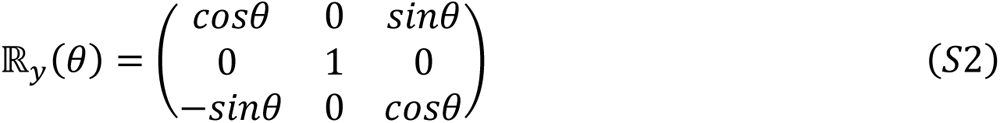

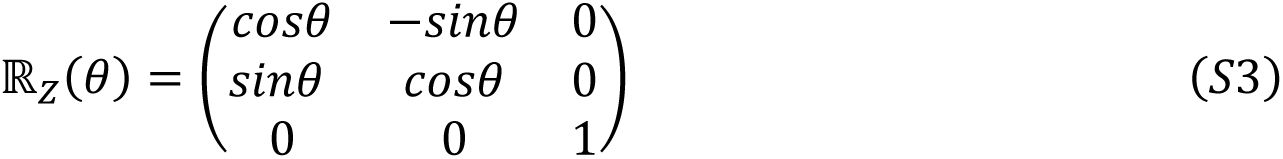

If 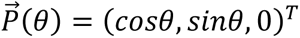 is the initial location of a 3-D point on the object lying on the XY plane with radius 1, the position of the point 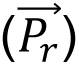 on a rotating ring inclined at an angle of *τ* from the ground plane and angular velocity of *ω* is expressed as:

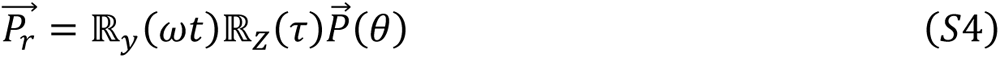

The position of the point on the wobbling object will be:

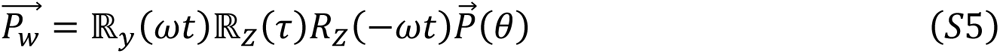

The operations of the rotational matrix for wobbling and rotation are illustrated in Figure 5 D. In Equation S10 (see below), 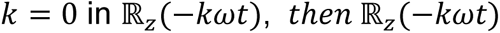 is the identity matrix. Thus, Equation S10 provides a smooth transition from rotation to wobbling. Then, we projected each point in perspective for the stimulus, but for the analysis, we used orthographic projection. For the physical stretch, 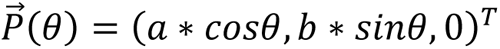 *a* and *b* were changed to make changes in aspect ratio that led to narrower or wider ellipses. Image stretches were performed on the projected image.

To compute velocity fields, the time derivative was taken as shown in Equation S11 (see below).

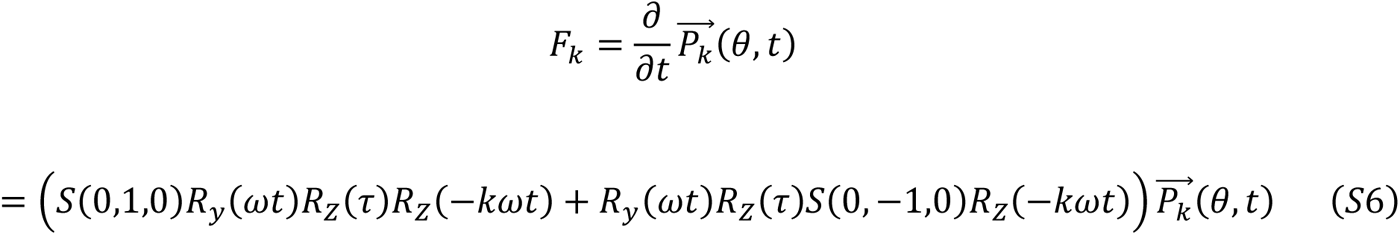

where *S*(0,1,0) is a skew symmetric matrix 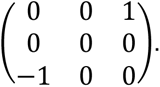

### DECODING SHAPE

We decode shapes from the responses of cortical neurons selective for orientation. In Figure 3 A& B, the simulated orientation tuning curves use von Mises probability density functions to generate the tuning curve for all orientations θ:

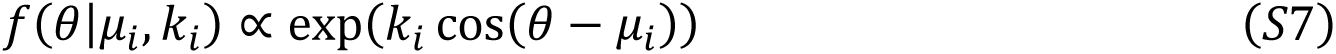

Where μ_*i*_ is the preferred orientation of the *i*-th cell and *k*_*i*_ sets the tuning width. Since we will be decoding angles, two orientations 180° apart are considered distinct, so Equation 1 covers 360°, unlike most fits of the von Mises function^36^. As a control, we also decoded angles from a cortex with isotropic distribution of orientation preference and tuning (Figure 3A).

#### Perceived angle decoded from cortical responses

We decoded the difference between angles around the horizontal axis (*γ*_*h*_) and angles around the vertical axis (*γ*_*v*_), for angles ranging from 10-170° from an anisotropic cortex and an isotropic cortex. We assume that each angle γ is formed by two lines θ_*a*_ and θ_*b*_, to which the response of *n*_*i*_ neurons with preferred orientation μ_*i*_ are *r*_*ai*_ and r_*bi*_ :

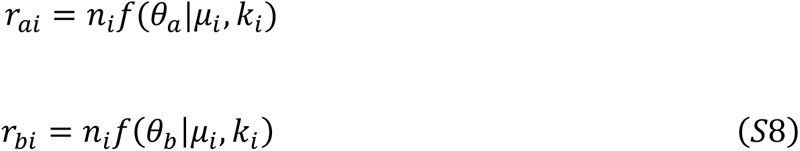

Each response is normalized by a divisive gain control:

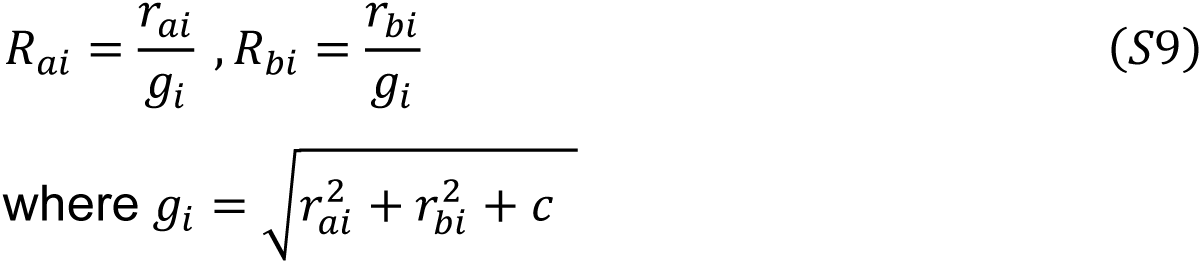

The line orientations, θ_*a*_ and θ_*b*_, are decoded using vector sums^41, 42^:

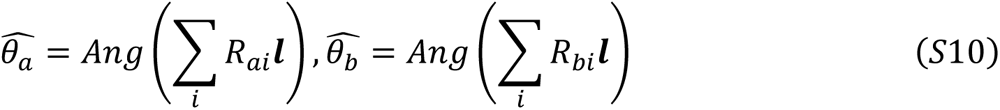

Where *l* is the unit vector that points to the preferred orientation. Then, the decoded angle is simply calculated from the two decoded line orientations:

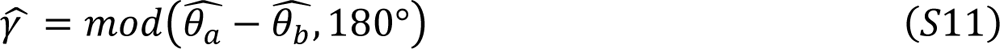

### DECODING NON-RIGIDITY VERSUS RIGIDITY

#### Nonrigidity anisotropy from cortical anisotropy in direction selectivity

To simulate the effect of spatial anisotropy on optic flow, we computed local motion energy for each pixel of the videos. Each motion-energy unit consisted of a quadrature pair of 3D Gabor filters ^48–55^:

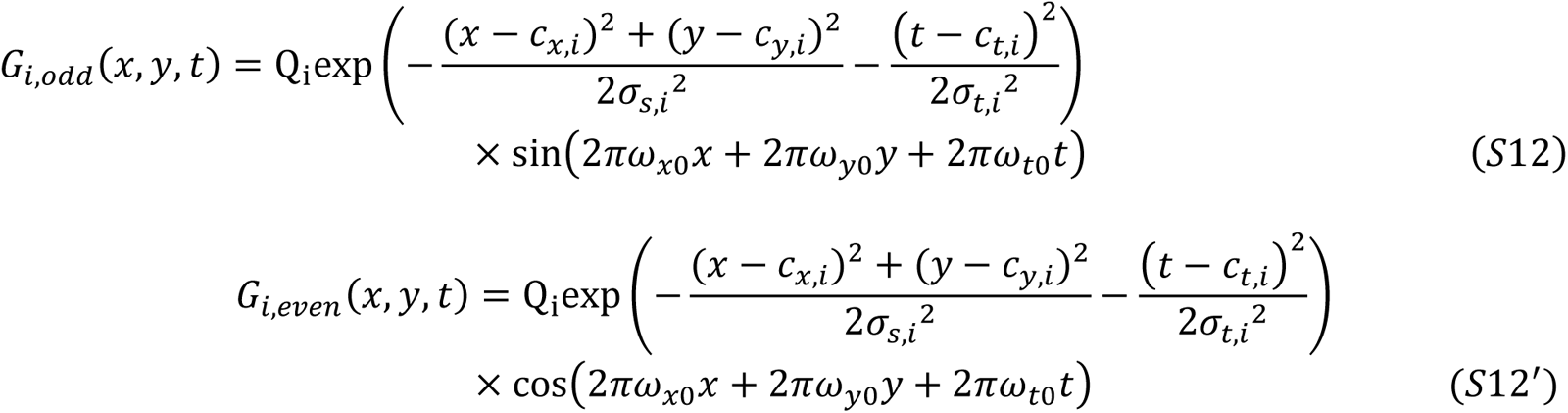

Where *Q*_*i*_ is the normalization constant; and *c*_*x,i*_ and *c*_*y,i*_ are the spatial means of the Gaussian envelopes, and *c*_*t,i*_ is the temporal mean; *σ*_*s,i*_ and *σ*_*t,i*_ are the spatial and temporal standard deviations of the Gaussian envelopes, respectively; and *ω*_*x*0_, *ω*_*y*0_, and *ω*_*t*0_ are the spatial and temporal frequencies of the sine component of the Gabor filter, referred to as the preferred spatial and temporal frequencies where:

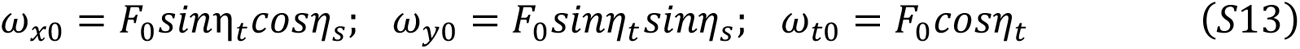

η_*s*_ and η_*t*_ are spatial and temporal orientations (8 spatial orientations from 0 to 7/8π and 5 temporal orientations from −π/4 to π/4) and σ_0_ is the frequency (0.28 and 0.14 cycles/pixel). For the anisotropic cortex, *σ* is adjusted so that the half-height orientation bandwidths 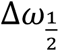 are matched to cat cortex estimates^21^ by the following equation^56^:

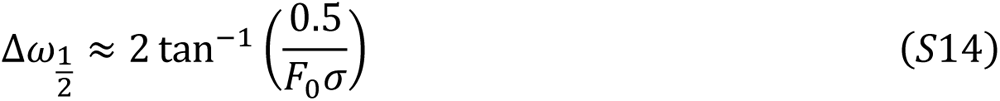

Each filter *G*_*i,even*_ and *G*_*i,odd*_ was convolved with the video I(*x*, *y*, *t*). The responses of the quadrature pair were squared and summed to produce a phase-independent response *m*_*i*_^48–55^. Figure 5 A and B illustrate the direction tuning curves for isotropic and anisotropic cortices, weighted by the number of cells. In the anisotropic cortex, cells that prefer a vertically moving bar (either upwards or downwards) exhibit narrower tuning and are more numerous compared to cells that prefer oblique or horizontally moving bars.

To obtain the best estimate of velocity 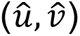 per point using the whole family of filters, we computed optic flow by estimating the motion vector (*u*, *v*) that minimizes the difference between the predicted and measured motion energies^57^:

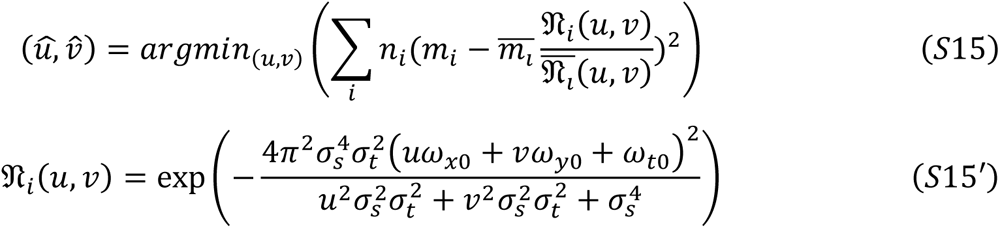

Where 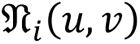 is the predicted response of filter *i* to velocity (*u*, *v*), *m*_*i*_ is the motion energy output, and 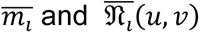 are the sum of 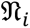 and the sum of *m*_*i*_ for filters sharing the same spatial orientation. Figure 5C shows optic flow fields for different conditions of rotation of a single ring, horizontal rotation ℝ_*y*_ around the y-axis (left column), and vertical rotation ℝ_*x*_ around the x-axis (right column), for the isotropic cortex (top row), and the anisotropic cortex (bottom row). The color of the vector corresponds to the direction and the length represents the magnitude of the estimated velocity. Most of the optic flow vectors from the isotropic cortex are orthogonal to the contour. In contrast, for the anisotropic cortex, the optic flow vectors shift towards the horizontal direction, as indicated by the red and turquoise colors.

To understand what this horizontal shift in the optic flow implies for perceived rotation versus wobble (which adds rotation ℝ_*z*_ around the z-axis), we derived the velocity field for different combinations of physical rotation and wobbling of a single ring, with the wobbling weight *k* ranging from *k* = 0∼*rotation* to *k* = 1∼*wobbling*. We began with the vector 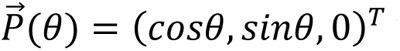 which defines the position of every point on a ring lying on the *X* − *Z* plane for each angular coordinate of the circular ring θ (Figure 5D), and calculated 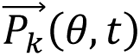 for every combination of rotation and wobble given by ℝ_*y*_ and ℝ_*z*_ the rotational matrices around the *Y* and *Z* axis respectively, with the angular velocity *ω* and the tilt of the ring, *τ*:

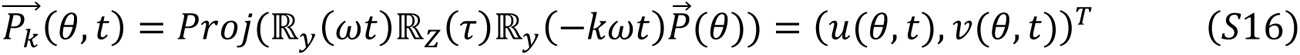

*Proj*(·) is a projection function and we used the orthographic projection. Then, the velocity field σ_*k*_ is calculated by the partial derivative with respect to time:

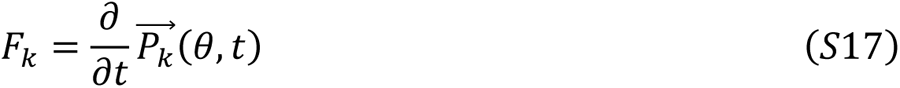

Figure 5E illustrates the velocity fields for horizontal rotation (top) and vertical (bottom), obtained with wobbling weights (*k*) increasing from left to right, from pure rotation to pure wobbling. For horizontal rotation (top row), horizontal vectors are predominant when *k* = 0 and vertical vectors at *k* = 1, with the opposite for the vertical rotation (bottom row). These velocity fields serve as templates, and the cosine similarity between these derived templates and the estimated optic flows in Figure 5C is calculated at each point and time for both the isotropic and anisotropic cortex (Figures 5F & 5G) and then averaged over all points for a complete cycle. The blue trace represents mean cosine similarity for horizontal rotation, and the red trace for vertical rotation. Having all possible templates is biologically implausible, so next we explore how a factoring of the velocity fields by more biologically plausible filters can lead to similar comparisons that predict the nonrigidity anisotropy.

#### Nonrigidity anisotropy from biologically plausible filters for differential invariants

In kinematics, which describes the motion of objects without considering the forces that cause them to move, motion fields are analyzed in terms of the local differential invariants, divergence, curl, and deformation after excluding translation parallel to the image plane, and these analyses have been useful in understanding object motion from optic flow^35, 58, 59, 60^. For the single ring with some combination of rotation and wobble, the divergence field is perpendicular to the contour and captures expansion/contraction of the projection (Figure 6A); the curl field is tangential to the contour and captures rotation; the deformation fields captures shear caused by the difference between orthogonal axes in contraction/expansion or shear and can be combined into one measure giving direction and magnitude of deformation^60^.

To derive divergence, curl, and deformation fields for physical rotation and physical wobbling, the dot product of the velocity fields derived in Equation S17 as a function of *k* (illustrated in Figure 5E) was taken with each operator corresponding to those illustrated in Figure 6A, and then integrated across space to obtain one number for each invariant per phase of motion, where ***i*** is the basis vector (1,0) and ***j*** is (0,1) :

Divergence:

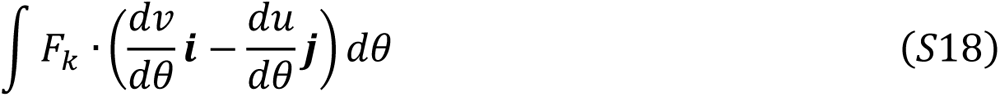

Curl:

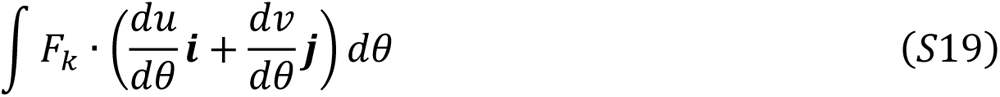

Deformation:

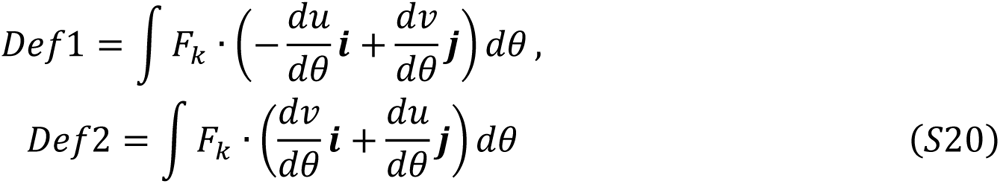

These are combined into one measure of the magnitude of Def:

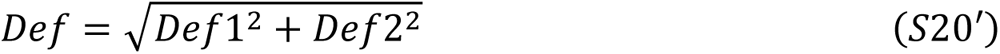

Figure 6B displays the values of derived integrated gradients for divergence, curl, and deformation from left to right as a function of the motion phase. The colors represent different wobbling weights (*k*), with warmer colors indicating more wobbling and cooler colors representing more rotation.

### Equation symbol lookup table

| Symbol | Description |
| --- | --- |
| $\phi_v, \phi_h, \psi_v, \psi_h$ | Angles in two elongated diamonds depicted in Figure 2C and D |
| $k_i, \mu_i$ | Kappa sets the tuning width and Mu the mean of the von Mises function for the i-th neuron |
| $\gamma_h, \gamma_v$ | Angles around the horizontal axis and the vertical axis |
| $\theta_a, \theta_b$ | Orientation of two lines that form an angle of $\gamma$ |
| $n_i$ | Number of cells for i-th neuron |
| $r_{ai}, r_{bi}$ | Feedforward response of i-th cell given orientations of $\theta_a$ and $\theta_b$ respectively |
| $g_i$ | Divisive gain control of i-th cell |
| $R_{ai}, R_{bi}$ | Normalized response of i-th cell |
| $\widehat{\theta}_a, \widehat{\theta}_b$ | Decoded orientations |
| $\widehat{\gamma}_v, \widehat{\gamma}_h$ | Decoded angles given $\gamma_h$ and $\gamma_v$ respectively |
| $G_{i,odd}, G_{i,even}$ | Odd and even 3D Gabor filters of i-th cell |
| $Q_i$ | Normalization constant for the Gabor filter |
| $c_{x,i}, c_{y,i}, c_{t,i}$ | Spatial and temporal means of the Gabor filter |
| $\sigma_{s,i}, \sigma_{t,i}$ | Spatial and temporal envelopes of the Gabor filter |
| $\omega_{x0}, \omega_{y0}, \omega_{t0}$ | Spatial and temporal frequencies of the sine component of the Gabor filter |
| $\eta_s, \eta_t$ | Spatial and temporal orientations of the Gabor filter |
| $F_0$ | Frequency of the Gabor filter |
| $\Delta\omega_{\frac{1}{2}}$ | Half-height orientation bandwidths |
| $\hat{u}, \hat{v}$ | The best estimate of local velocity components |
| $\mathfrak{N}_i(u, v)$ | Predicted response of filter $i$ to velocity $(u, v)$ |
| $m_i$ | $i$-th motion energy output |
| $\overline{m}_i, \overline{\mathfrak{N}}_i(u, v)$ | The sum of $\mathfrak{N}_i$ and the sum of $m_i$ for filters sharing the same spatial orientation |
| $\mathbb{R}_y, \mathbb{R}_z$ | Rotational matrices around Y and Z axis respectively |
| $\vec{P}(\theta)$ | The position of a point on the ring lying on the $X - Z$ plane |
| $\tau$ | Tilt of the ring |
| $\omega$ | Angular velocity of the rotation of the ring |
| $k$ | The wobbling weight ($k = 0 \sim \text{rotation}, k = 1 \sim \text{wobbling}$) |
| $Proj(\cdot)$ | The projection function |
| $\vec{P}_k(\theta, t)$ | The position of a point at a space and time for a moving ring with the wobbling weight of $k$ |
| $u(\theta, t), v(\theta, t)$ | The $x$ and $y$ components for $\vec{P}_k(\theta, t)$ |
| $F_k$ | The velocity field for the moving ring with the wobbling weight of $k$ |

**Figure S1.**
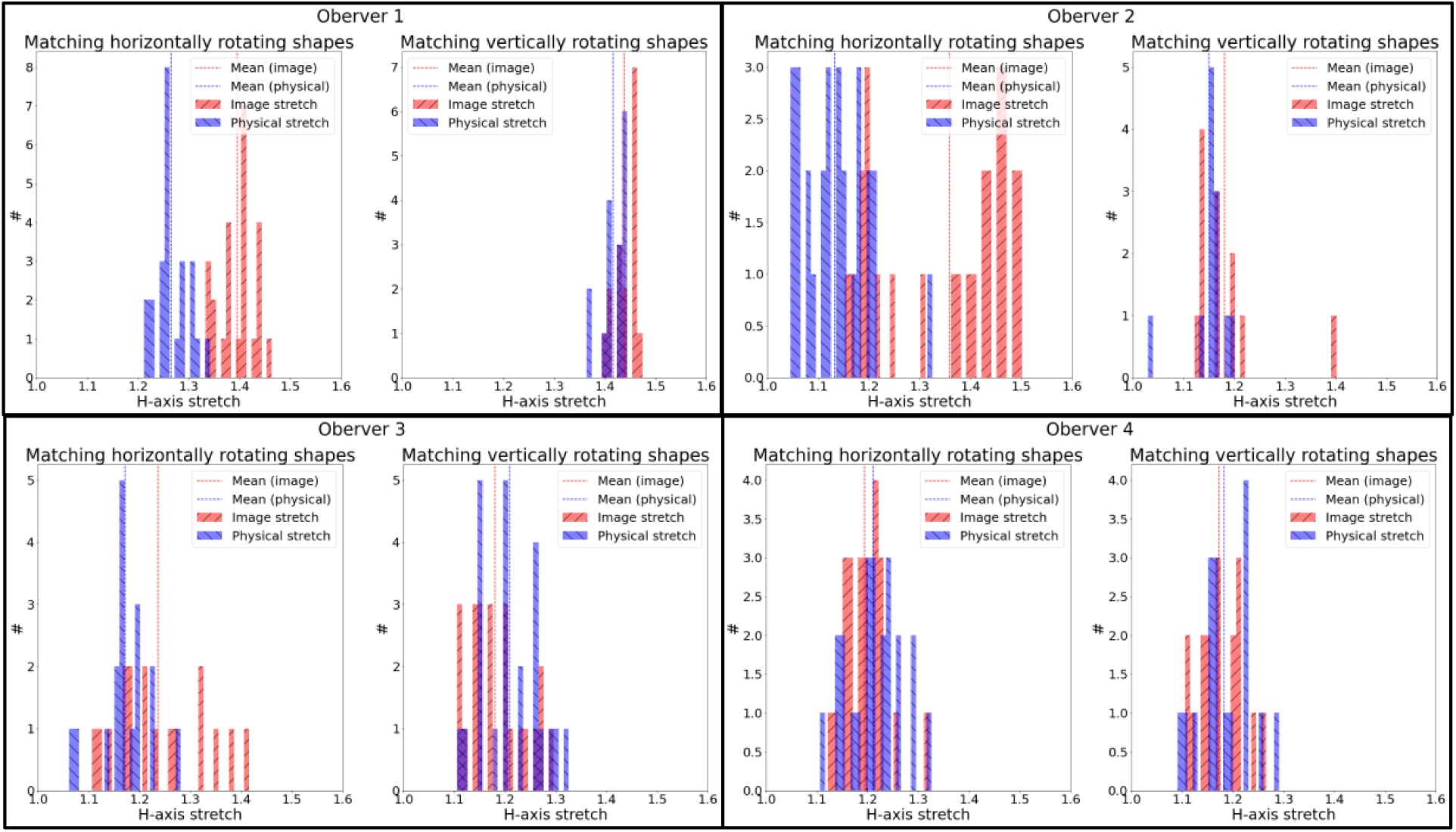
Individual results for the shape matching experiment. For each observer, left histograms represent the extent of horizontal stretch in the image domain (red) and physical domain (blue) for vertically rotating rings adjusted to match the shape of physically circular horizontally rotating rings. Right histograms represent the extent of horizontal stretch in the image domain (red) and physical domain (blue) for horizontally rotating rings adjusted to match the shape of physically circular vertically rotating rings.

**Figure S2.**
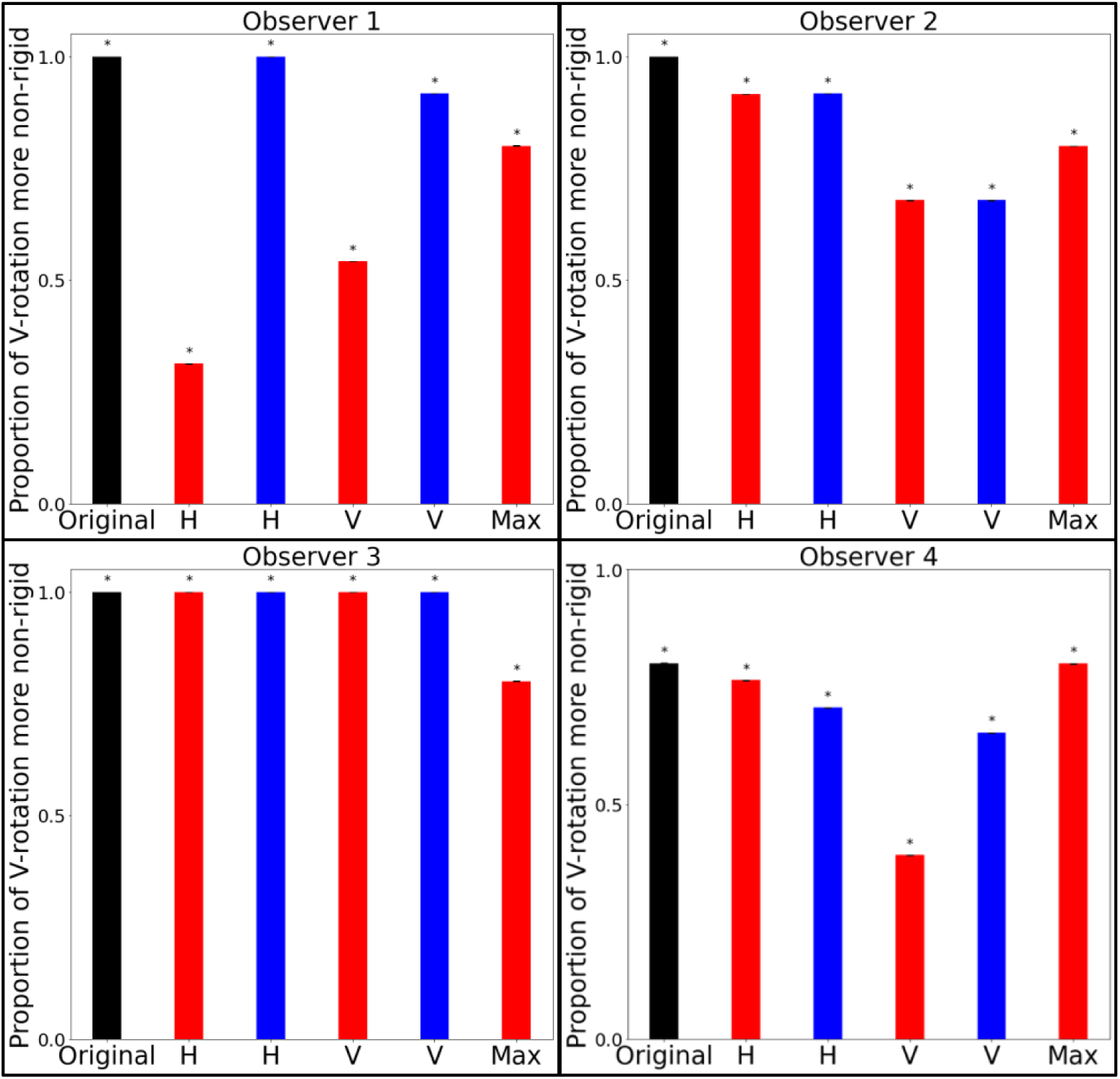
Individual results for the nonrigidity comparison. Probability of each observer reporting vertically rotating rings as more nonrigid. Original: Two pairs of circular rings (black). H indicates that horizontally rotating rings were stretched to match shapes, and V indicates that vertically rotating rings were stretched, either in the image (red) or physically before projection (blue). Max: Both vertical and horizontal rings were stretched maximally like in the last column of Panel A in Figure 4, so horizontally rotating rings were perceived as narrower and longer than vertically rotating rings.

